# Individual susceptibility to TMS affirms the precuneual role in metamemory upon recollection

**DOI:** 10.1101/383299

**Authors:** Qun Ye, Futing Zou, Michael Dayan, Hakwan Lau, Yi Hu, Sze Chai Kwok

**Author notes:** Correspondence (Sze Chai Kwok), Room 269, Geography Building, 3663 Zhongshan Road North, Shanghai 200062, China.

## Abstract

**Background:** A recent virtual-lesion study using inhibitory repetitive transcranial magnetic stimulation (rTMS) confirmed the causal behavioral relevance of the precuneus in the evaluation of one’s own memory performance (aka mnemonic metacognition).

**Objective:** This study’s goal is to elucidate how these TMS-induced neuromodulatory effects might relate to the neural correlates and be modulated by individual anatomical profiles in relation to meta-memory.

**Methods:** In a within-subjects design, we assessed the impact of 20-min rTMS over the precuneus, compared to the vertex, across three magnetic resonance imaging (MRI) neuro-profiles on 18 healthy subjects during a memory versus a perceptual task.

**Results:** Task-based functional MRI revealed that BOLD signal magnitude in the precuneus is associated with variation in individual meta-memory efficiency, and such correlation diminished significantly following TMS targeted at the precuneus. Moreover, individuals with higher resting-state functional connectivity (rs-fcMRI) between the precuneus and the hippocampus, or smaller grey matter volume in the stimulated precuneal region exhibit considerably higher vulnerability to the TMS effect. These effects were not observed in the perceptual domain.

**Conclusion:** We provide compelling evidence in outlining a possible circuit encompassing the precuneus and its mnemonic midbrain neighbor the hippocampus at the service of realizing our meta-awareness during memory recollection of episodic details.

**Highlights:**

- TMS on precuneus reduces meta-memory ability during memory retrieval.
- TMS disrupts the correlation between BOLD activity and meta-memory ability.
- TMS effect is modulated by rs-fcMRI between precuneus and hippocampus.
- Individuals with greater precuneal grey matter volume more immune to TMS effect.

## Introduction

The ability to accurately monitor and evaluate one’s own behavioral performance is a critical feature of our cognitive function. Recent studies have advanced our understanding of the neural underpinnings of metacognitive ability, mainly with a focus on the perception and memory domains. While ample neuroimaging and neuropsychological evidence from distinct modalities convergently point to the anterior prefrontal cortex (aPFC) being specifically related to perceptual metacognition, including white matter (WM) fiber tracking [1], microstructural measures of WM concentration [2], grey matter (GM) volume [1, 3], task-related functional magnetic resonance imaging (fMRI) [4-7], resting-state fMRI [8], neurophysiology [9, 10], and lesion-based studies [11-13], our understanding of the neural correlates of metacognition for memory is in contrast less conclusive.

Researchers have used a combination of objective memory task accuracy (usually from recognition or forced-choice tasks, known as type 1 tasks) and subsequent subjective confidence rating (type 2 tasks) to define successful decision making [14], and have found the memory-related signals and the confidence-related signals can diverge and might rely on two largely independent processes [15]. Neurally, other investigations have implicated the posterior parietal cortex in the subjective experiences and mnemonic metacognition of memory contents [16]. Most notably, patients with lesions on the posterior parietal cortex tend to show less confidence in their source recollection even though their type 1 task appears to be executed as well as healthy controls [16, 17], implying a critical role of the parietal cortex in mnemonic metacognition [18]. It has also been shown that the medial parietal cortex was particularly activated during confidence rating in memory tasks [6, 19] and that individual differences in mnemonic metacognition ability to be correlated with resting-state connectivity between the aPFC and the right precuneus [8], as well as with variation in the volume of the precuneus [3].

In a previous paper [18], we have confirmed the causal relevance of the precuneus in mnemonic metacognition via inhibiting the normal functioning of the precuneus temporarily with non-invasive low-frequency repetitive transcranial magnetic stimulation (1-Hz rTMS). However, we have not been able to characterize the individual neural variability affected by the neuromodulatory effects of the TMS on this critical region for this metacognitive process. Hence, here we tried to capitalize on the individual neural variability in combination of TMS to elucidate the neural correlates of meta-memory using both functional and structural data.

Our experimental protocol utilized data from subjects’ resting-state functional MRI, structural MRI, and two task-based functional MRI following stimulation on a target region, the precuneus, or a control region, the vertex. We analyzed the resting-state functional connectivity, voxel-based morphometry (VBM), and blood oxygen level dependent (BOLD) activity by specifically comparing the putative effects of TMS on the precuneus with a control stimulation condition. Our results provided a comprehensive profile to characterize the neuromodulatory effects by the focal magnetic stimulation on the precuneus across subjects through the three different MRI analyses, and corroborated previous findings on the contribution made by the precuneus in supporting mnemonic metacognition [3, 18].

## Material and methods

### Participants

Participants were recruited from the student community of the East China Normal University and were compensated for their participation. Data came from 18 healthy adults (7 females; age 19-24 years). Each of them participated in two experiments, each including two TMS sessions, giving us a within-subjects comparison. No subjects withdrew due to complications from the TMS procedures, and no negative treatment responses were observed. The number of participants was decided based on previous work adopting a similar experimental design [20]. All participants had normal or corrected-to-normal vision, no reported history of neurological disease, no other contraindications for MRI or TMS, and all gave written informed consent. The study was approved by University Committee on Human Research Protection of East China Normal University (UCHRP-ECNU).

### Overview of study

Participants initially underwent a high-resolution structural MRI scan, which was used to define subject-specific coordinates for subsequent sites of TMS and voxel-based morphometry analysis and undertook a 7 min of resting-state fMRI scan, which was used for subsequent functional connectivity analysis. The participants were asked to complete two experimental sessions of a memory task. For each session of the memory part, participants needed to play one video game (encoding phase, out of MRI scanner), and received 20 min of rTMS (rTMS stimulation phase) before going to complete the memory retrieval task conducted in MRI scanner (memory retrieval phase). The chapters of the video game and the stimulation sites were counterbalanced across the two sessions (Figure 1A). The same participants also participated in a perceptual experiment outside the scanner for task comparison purpose.

**Figure 1.**
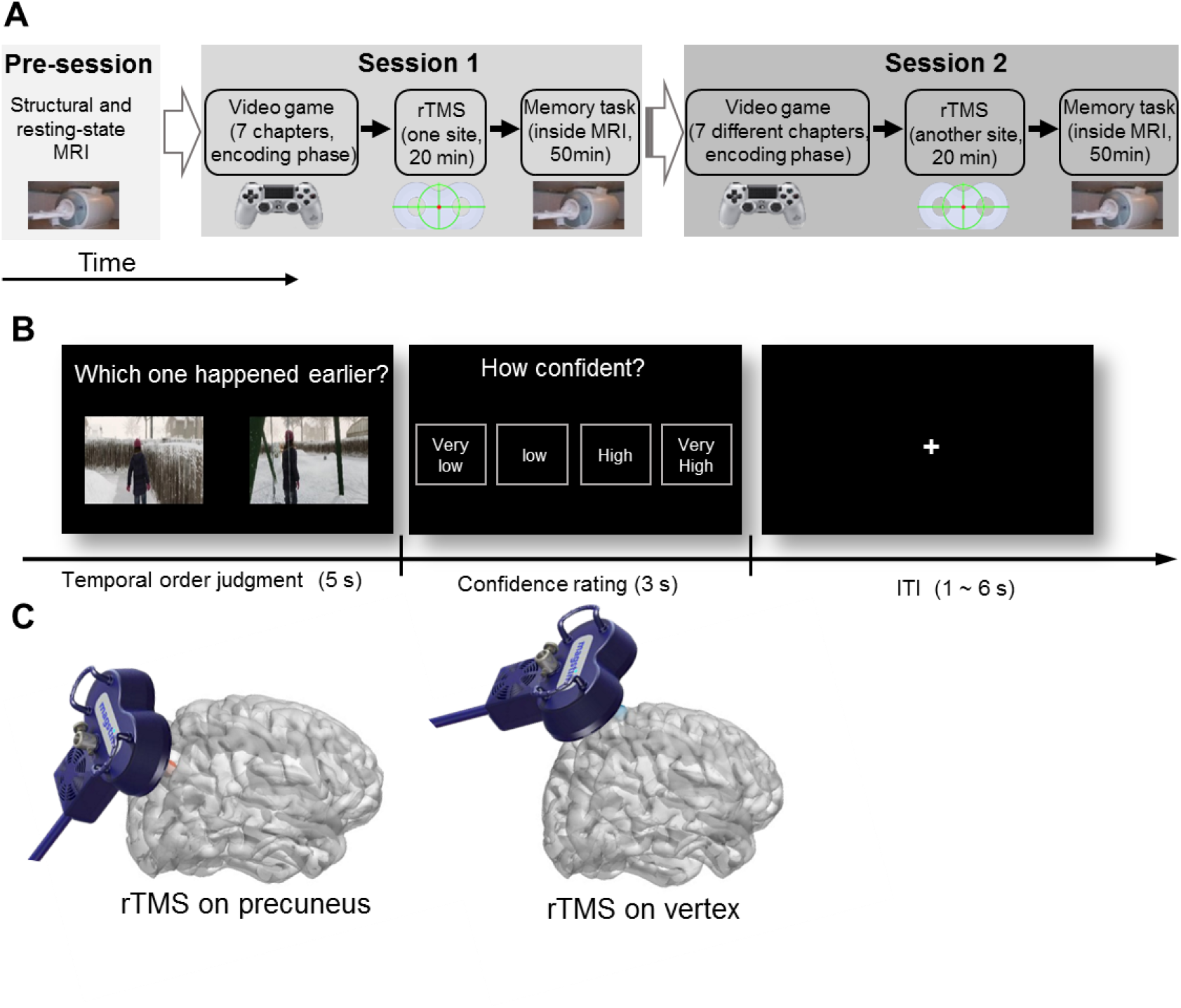
(A) Experiment overview. Participants underwent structural and resting-state MRI scans during the pre-session. In session 1 and session 2, participants played a video game containing seven related chapters, and 24 hours later, received 20 min of repetitive transcranial magnetic stimulation (rTMS) to either one of two cortical sites before performing a memory retrieval task during MRI. The two sessions were conducted within-subjects on two different days. (B) Temporal order memory retrieval task. Participants chose the image that happened earlier in the videoplay they had played and reported their confidence rating on how confident their judgment was correct, from very low to very high. (C) TMS to stimulation sites. Location of precuneus (target site, MNI coordinates: x, y, z = 6, −70, 44) is depicted with a red dot (left) and vertex with a blue dot (right).

**Figure 2.**
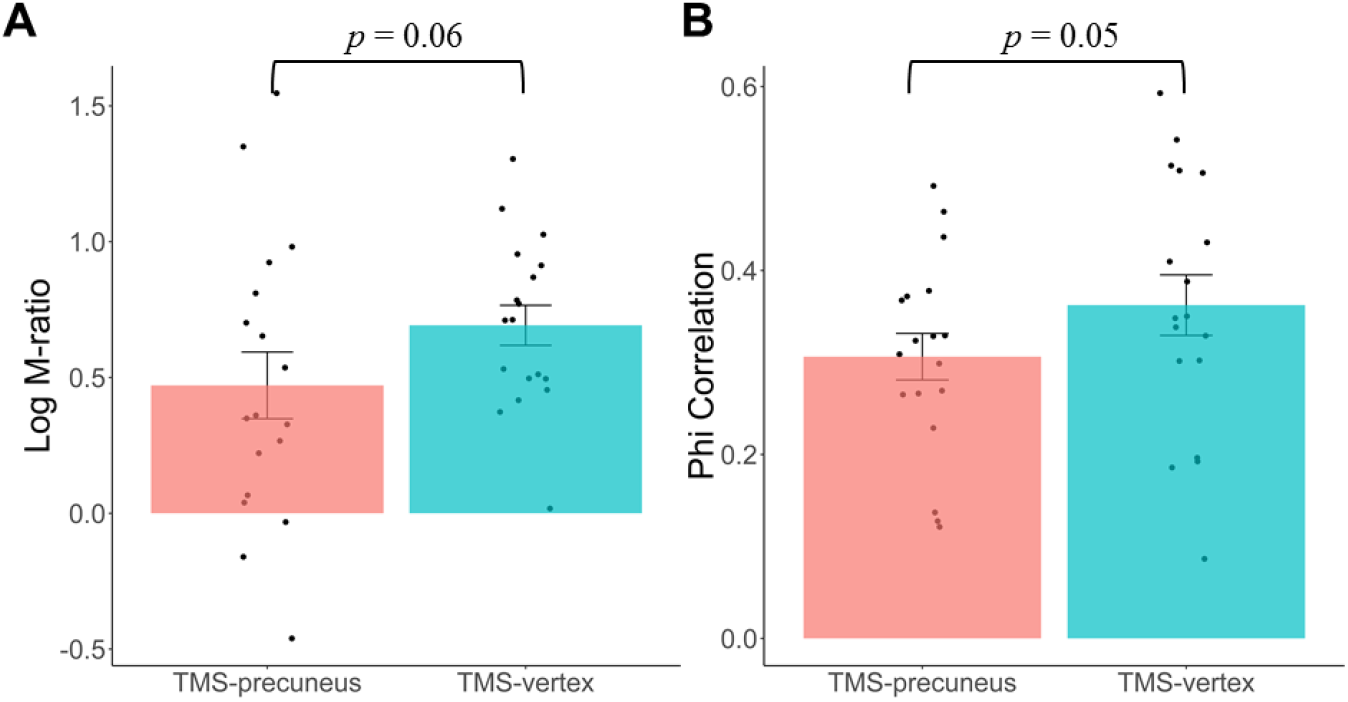
Effects of TMS on meta-memory efficiency. Metacognitive ability was reduced after TMS to precuneus compared to TMS to vertex in (A) SDT metacognitive efficiency measure (Log M-ratio) and (B) Phi coefficient. Black dots denote metacognitive score per subject. Error bars denote the standard error of the mean (SEM) over participants.

### Stimuli

The stimuli were extracted from an action-adventure video game (*Beyond: Two Souls*, Quantic Dream, France; PlayStation 4 version, Sony Computer Entertainment.). The Participants played 14 chapters in total across two sessions: 7 in session 1 and then another 7 in session 2. These subject-specific videos were recorded and were used for extraction of still images in both sessions.

For the memory task, we selected still frames/images from the subject-specific recorded videos in which participants had played the day before. Each second in the video consisted of 29.97 static images. In each game-playing session, 240 pairs of images were extracted from the 7 chapters and were paired up for the task based on the following criteria: (1) the two images had to be extracted from either the same chapters or adjacent chapters (Within-vs. Across-chapter condition); (2) the temporal distance (TD) between the two images were matched between Within- and Across-chapter condition; (3) in order to maximize the range of TD, we first selected the second longest chapter of the video and determined the longest TD according to a power function (power = 1.5). We generated 60 progressive levels of TD among these pairs. For the perceptual task, we used the same set of subject-specific stimuli generated for the memory task to rule out any potential stimuli idiosyncrasy. The resolution of one of the paired images was changed using Python Imaging Library by resizing the image to modulate the pixel dimension; this modification of image resolution was conducted for an image-resolution comparison task (see below).

### TMS: sites, protocol, and procedure

Repetitive transcranial magnetic stimulation (rTMS) was applied using a Magstim Rapid^2^magnetic stimulator connected with a 70mm double air film coil (The Magstim Company, Ltd., Whitland, UK). The subject-specific structural T1 images were obtained and used in the Brainsight2.0 (Rogue Research Inc., Montreal, Canada), a neuronavigation system, coupled with infrared camera using a Polaris Optical Tracking System (Northern Digital, Waterloo, Canada), to localize the target brain sites. Target stimulation sites were selected in the system by transformation of the Montreal Neurological Institute (MNI) coordinates to participant’s native brain. The stimulation sites located in the precuneus at the MNI coordinate x=6, y=-70, z=44 [21], and in a control area on the vertex, which was identified at the point of the same distance to the left and the right pre-auricular, and of the same distance to the nasion and the inion (Figure 1C). To prepare the subject-image registration and promote on-line processing of the neuronavigation system, four location information of each subject’s head were obtained manually by touching fiducial points, which are the tip of the nose, the nasion, and the inter-tragal notch of each ear using an infrared pointer.

In each session, rTMS was delivered to either the precuneus or vertex site before the participants engaged in performing the memory/perceptual tasks. rTMS was applied at low-frequency for a continuous duration of 20 min (1 Hz, 1,200 pulses in total) at 110% of active motor threshold (MT), which was defined as the lowest TMS intensity delivered over the motor cortex necessary to elicit visible twitches of the right index finger in at least 5 out of 10 consecutive pulses. The MT was measured prior to administering the stimulation (MT range: 57% - 80%; mean ± sd: 68.28% ± 6.19%). During stimulation, participants wore earplugs to attenuate the sound of the stimulating coil discharge. The coil was held to the scalp of the participant with a custom coil holder and the subject’s head was propped with a comfortable position. Coil orientation was parallel to the midline with the handle pointing downward. Immediately after the 20 min of rTMS, subjects performed four blocks of memory retrieval task inside MRI scanner. This particular stimulation magnitude and protocols of rTMS (low-frequency stimulation of 1 Hz) is known to induce efficacious intracortical inhibitory effects for over 60 min [22, 23]. Given that each session of the memory/perceptual tasks lasted approximately 45 min, the TMS effects should have been long-lasting enough for the tasks. For safety reason and to avoid carry-over effects of rTMS across sessions, session 1 and 2 of both tasks were conducted on two separate days.

### Memory task (temporal-order judgment, TOJ), perceptual task (image-resolution judgment), and confidence ratings

The memory retrieval task required participants to choose the image that happened earlier in the video game they had played one day before (temporal order judgment, TOJ). The memory retrieval task was administrated inside an MRI scanner, where visual stimuli were presented using E-prime 2.0 software (Psychology Software Tools, Inc., Pittsburgh, PA), as back-projected via a mirror system to the participant. Each trial started by a temporal order judgment in 5 s, and immediately followed by a confidence judgment within 3 s. Participants performed the temporal order judgment using their index and middle fingers of one of their hands via an MRI compatible five-button response keyboard (Sinorad, Shenzhen, China), and reported their confidence level (“Very Low”, “Low”, “High”, or “Very High”) regarding their own judgment of the correctness of the TOJ with four fingers (thumb was not used) of the other hand. The left/right hand response contingency was counterbalanced across participants. Participants were encouraged to report their confidence level in a relative way and make use of the whole confidence scale. Confidence judgments are one commonly used method for quantifying the sensitivity of self-reported confidence to objective discrimination performance under the signal detection theory [24]. These confidence ratings will be used in our computation for metacognitive indices (see below). Following these judgments, a fixation cross with a variable duration (1 –6 s) was presented (Figure 1B). For either of the sessions,there was a practice block for participants to get familiar with the task before going into MRI scanner. In total, each participant completed 240 trials in either of the sessions (4 blocks × 60 trials).

The perceptual task required participants to choose either the clearer (or blurrier, counter-balanced across participants) image among a pair of images on each trial. An identical confidence rating procedure as of the memory task was adopted immediately following each image-resolution comparison judgment. Each participant completed 240 perceptual discrimination trials in each of the two sessions.

### MRI data acquisition

All the participants were scanned in a 3-Tesla Siemens Trio magnetic resonance imaging scanner using a 32-channel head coil (Siemens Medical Solutions, Erlangen, Germany). A total of 1,350 fMRI volumes and 220 rs-fMRI volumes were acquired for each subject. The functional images were acquired with the following sequence: TR = 2000 ms, TE = 30 ms, field of view (FOV) = 230 × 230 mm, flip angle = 70°, voxel size = 3.6 × 3.6 × 4 mm, 33 slices, scan orientation parallel to AC-PC plane. High-resolution T1-weighted MPRAGE anatomy images were also acquired (TR = 2530 ms, TE = 2.34 ms, TI = 1100 ms, flip angle = 7°, FOV = 256 × 256 mm, 192 sagittal slices, 0.9 mm thickness, voxel size = 1 × 1 × 1 mm).

### Data analysis

#### Behavioral data analysis

We evaluated the metacognitive ability by Meta-d’ using both memory performance and confidence ratings data. Meta-d’ quantifies metacognitive sensitivity (the ability to discriminate between one’s own correct and incorrect judgments) in a signal detection theory (SDT) framework. Meta-d’ is widely used as a measure of metacognitive capacity and expressed in the same units as d’, so the type 2 sensitivity (meta-d’) can be compared with the type 1 sensitivity (d’) directly [14, 24]. If meta-d’ equals to d’, the participant makes confidence rating with maximum possible metacognitive sensitivity. If meta-d’ less than d’, the participant’s metacognitive sensitivity is suboptimal. Here, we calculated the logarithm of the ratio meta-d’/d’ (log M-ratio) for estimating the metacognitive efficiency (the level of metacognitive sensitivity given a particular level of performance capacity). The toolbox for the SDT-based meta-d’ estimation was available at http://www.columbia.edu/~bsm2105/type2sdt/.

In order to ensure our results were not due to any idiosyncratic violation of the assumptions of SDT, we additionally calculated the phi coefficient index, which does not make these parametric assumptions [14]. Rather, it evaluates how roughly “advantageously” each trial was assigned for high or low confidence based on performance in the preceding cognitive judgment, reflecting the association between the two binary variables [25]. The coefficient was calculated by the following equation using the number of trials classified in each case [n(case)]:

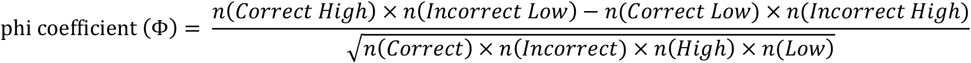

Trials missing either one of the measures (memory: 2.9% of TOJ trials, 2.2% confidence rating; perception: 0.7% of perceptual trials) were excluded from the analyses. The 4-point confidence ratings were collapsed into two categories (high vs. low) for analyses.

#### Task-based fMRI data analysis

Preprocessing was conducted using SPM12 (http://www.fil.ion.ac.uk/spm). Scans were realigned to the middle EPI image. The structural image was co-registered to the mean functional image, and the parameters from the segmentation of the structural image were used to normalize the functional images that were resampled to 3 × 3 × 3 mm. The realigned normalized images were then smoothed with a Gaussian kernel of 8-mm full-width half maximum (FWHM) to conform to the assumptions of random field theory and improve sensitivity for group analyses. Data were analyzed using general linear models as described below with a high-pass filter cutoff of 256 s and autoregressive AR(1) model correction for auto-correlation.

To identify brain areas in processing metacognitive information, we performed a contrast [(Correct_High – Correct_Low) > (Incorrect_High –Incorrect_Low)] at onsets of the memory phase with a duration of 5 s at single-subject level, including the following regressors: memory conditions (Correct, Incorrect, Miss) × confidence rating conditions (High, Low, Miss). Each run consisted of 6 head realignment parameters and the run mean were included as parameters of no interest. These events were modeled with a canonical hemodynamic response function as an event-related response. To test the relationship between the BOLD response and the behavioral meta-memory index (log M-ratio) across subjects, single-subject contrast images were entered into a second-level random effects analysis using one-sample *t* tests with log M-ratio as a covariate separately for two TMS sessions (TMS-precuneus vs. TMS-vertex).

The activation clusters were defined by the peak voxels on the normalized structural images and labeled using the nomenclature of Talairach and Tournoux (1988) [26]. Only activation surviving multiple correction at the cluster-level FWE corrected *p* < 0.005 threshold are reported below.

#### Resting-state functional connectivity analysis

A functional brain network was defined by a symmetric functional connectivity matrix c=c(i,j), where each row(i)/column(j) of the matrix is a network node, and each matrix entry c(i,j) is the weight of the network edge between node i and j. The connectivity matrices were obtained through a series of preprocessing steps on both rs-fMRI and T1 data, implemented in Python using a combination of fmriprep [27], nipype [28] and networkx packages (https://networkx.github.io/).

T1 preprocessing consisted in correcting for bias field using N4 [29], skull-stripping with ANTs (http://stnava.github.io/ANTs/), tissue segmentation into WM, GM and cerebral spinal fluid (CSF) with FSL (https://fsl.fmrib.ox.ac.uk/fsl/fslwiki) FAST, and non-linear registration to MNI space with ANTs. FreeSurfer (https://surfer.nmr.mgh.harvard.edu/) was used to reconstruct the GM and WM surfaces of each subject using the brain mask previously calculated, and to parcellate the brain into 86 regions as per the Desikan-Killiany atlas.

Resting-state preprocessing consisted in slice-time corrections with AFNI (https://afni.nimh.nih.gov/) 3dTShift, motion corrections with FSL MCFLIRT, and registration to the subject native T1 volume with FreeSurfer boundary based registration (using 9 degrees of freedom). ICA-based Automatic Removal of Motion Artifacts (AROMA) [30] was then applied to estimate noise regressors, while physiological noise regressors were calculated from voxels in the WM and CSF masks computed previously. The data was smoothed with an 8-mm kernel excluding background voxels with FSL Susan toolbox and all the noise components regressed out using FSL regfilt. Finally, bandpass filtering between 0.008 and 0.08 Hz was implemented with AFNI. Linear detrending was included in the previous step by adding a linear sequence as additional regressor. The FreeSurfer atlas was resampled to resting-state resolution, and the connectivity matrix calculated from the correlation of the denoised signal between each pair of atlas regions. Finally, the connectivity matrix entries were Fisher transformed. To remove spurious edges while ensuring consistent edge density across subjects, a lenient wiring cost of 50% was applied to all connectivity matrices which thus had half the total number of possible edges.

The edge length l(i,j) between two nodes was defined as the absolute inverse of the associated weight c(i,j), so that strong (i.e., high) correlation corresponded to short (i.e., low) length. We investigated in each hemisphere the hippocampus-precuneus (HP) connectivity distance which was defined as the shortest path length between these two nodes (computed using Dijkstra’s method) [31], that is the smallest sum of edge lengths among all the possible paths connecting them. The resulting HP connectivity distance was averaged across hemispheres.

#### Voxel-based morphometry (VBM) analysis

VBM preprocessing was performed using SPM12 (http://www.fil.ion.ucl.ac.uk/spm). Following the similar protocol used in previous studies [1, 3], the structural images were first segmented into GM, WM and CSF in native space. For increasing the accuracy of inter-subject alignment, the GM images were aligned and wrapped to an iteratively improved template using DARTEL algorithm, while simultaneously aligning the WM images [32]. The DARTEL template was then normalized to MNI stereotactic space, and then GM images were modulated in a way that their local tissue volumes were preserved. Finally, images were smoothed using an 8 mm full-width at half maximum isotropic Gaussian kernel.

The pre-processed images were analyzed in a multiple regression model to examine the relation between GM volume and difference in metacognitive efficiency between two TMS sessions (TMS-precuneus > TMS-vertex). Proportional scaling was used to account for volume variability in total intracranial volume across participants. A binary GM mask (> 0.3) was used to exclude clusters outside the brain and limit the search volume to voxels likely to contain GM.

We examined the positive and negative t-maps separately and identified clusters using an uncorrected threshold of *p* < 0.001 at voxel-level. These clusters were used to define regions of interest using MarsBar version 0.44 software (http://marsbar.sourceforge.net/). Following McCurdy et al. (2013)’s protocol [3], small-volume correction (SVC) was applied on a cluster of interest by centering a 10-mm sphere over the targeted site of stimulation in the precuneus (MNI: x=6, y=-70, z=44).

## Results

### Behavioral results

We first tested the hypothesis that TMS to the precuneus would reduce individual metacognitive ability in the memory task. There was a trend reduction in individual metacognitive efficiency in the TMS-precuneus session compared to the TMS-vertex session (Log M-ratio: paired t-test *t* (17) = 1.63, one-tailed*p* = 0.061). The trend was replicated with a SDT assumption free correlation measure computed by the association between the task performances and subsequent confidence ratings (*Phi* correlation: *t* (17) = 1.68, one-tailed *p* =0.055), and was confirmed by using another metacognitive efficiency measure, Meta-d’ –d’, in our previous paper [18]. Moreover, we ascertained that there were no significant differences in task performance and levels of confidence rating between the two TMS sessions (accuracy: paired t-test *t* (17) = 0.349, *p* = 0.640; confidence rating: *t* (17) = 0.070, *p* = 0.780). These results indicate that TMS to the precuneus specifically affected the individual metacognitive ability, and that no detectable effect related to their basic memory performance could be found.

### Task-based fMRI analysis

To test the effect of TMS on task-based BOLD responses, we correlated the interaction term [(Correct_High –Correct_Low) > (Incorrect_High –Incorrect_Low)] (i.e., difference in activation between correct vs. incorrect trials under high vs. low confidence) with log M-ratio index across subjects separately for TMS-vertex and TMS-precuneus sessions and compared the BOLD level of the interaction term between the two sessions. In the TMS-vertex session, there was a significant positive correlation between metacognitive efficiency and brain activation in one posterior cluster (k = 327 voxels, Figure 3A), extending from the precuneus (peak voxel, x, y, z = 6, −48, 14) to the posterior cingulate cortex (peak voxel, x, y, z = −3, −42, 14). No significant correlation between metacognitive efficiency and brain activation was found in the TMS-precuneus session. Note that there was no difference in the overall BOLD activation level indicated by the interaction term between the two TMS sessions (paired t-test *t* (17) = 0.44, *p* = 0.667). However, to further unpack these results, the activation cluster in TMS-vertex session was saved as a mask and we plotted the relationship between metacognitive efficiency and BOLD response separately for TMS-vertex and TMS-precuneus sessions. While the metacognitive efficiency was significantly correlated with the BOLD response in the TMS-vertex session (Pearson’s *r* = 0.76, *p* < 0.001), such correlational pattern was not observed in the TMS-precuneus session any more (Pearson’s *r* = 0.18, *p* = 0.475). The correlation coefficient was significantly lower than that of the TMS-vertex session (comparison between correlations: *z* = 2.36, *p* = 0.019; Figure 3B).

**Figure 3.**
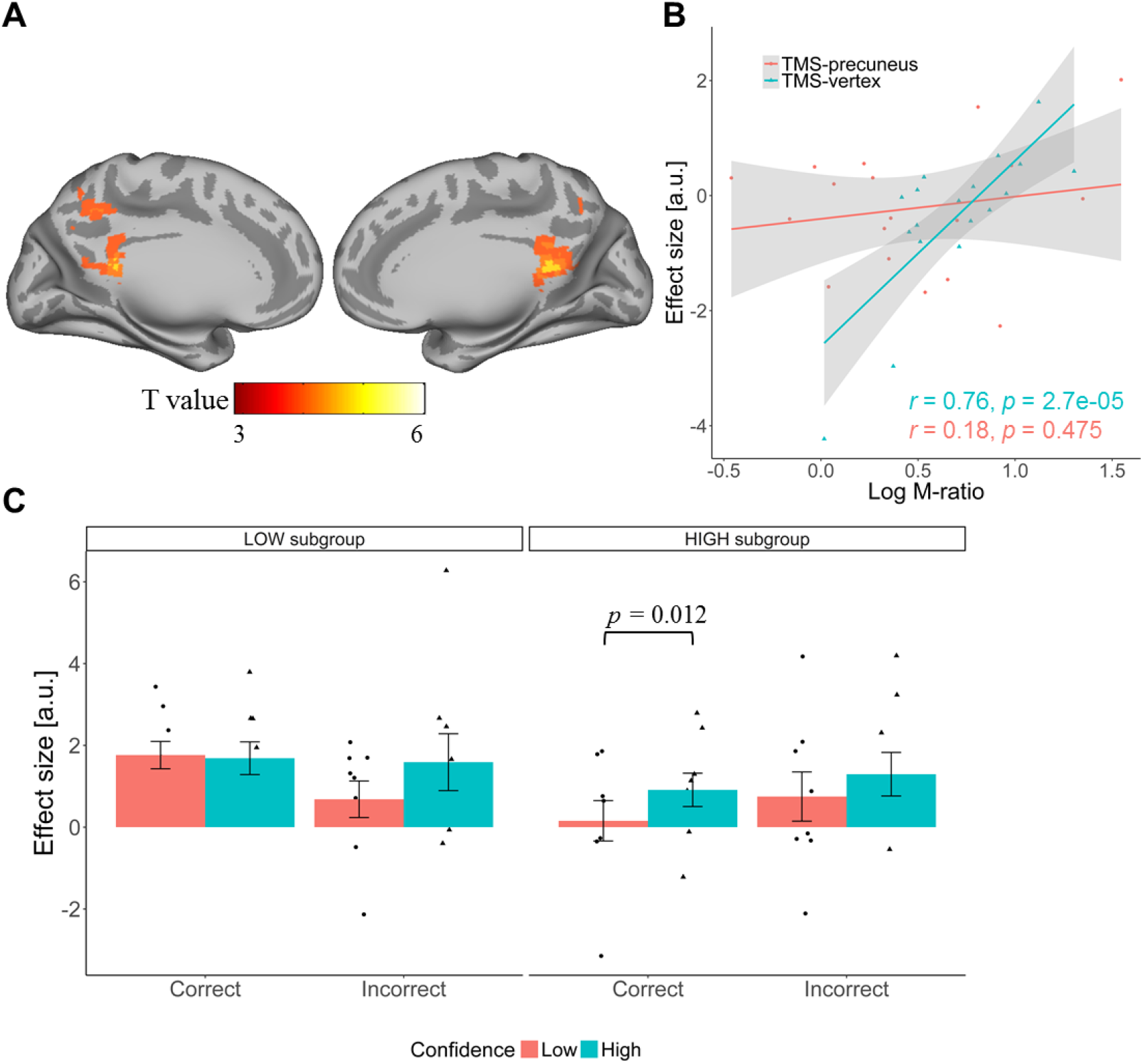
(A) Significant positive correlation between meta-memory efficiency (Log M-ratio) and activation in posterior medial region (peak voxel in precuneus: x, y, z = 6, −48, 14) in the TMS-vertex session. For visualization purposes, the threshold was set at voxel-level *p* < 0.005 uncorrected. (B) Individuals activation level (arbitrary unit, a.u.) in the precuneal cluster is correlated with meta-memory efficiency (Log M-ratio) only in the TMS-vertex session (cyan line) but not in the TMS-precuneus session (red line). Grey regions indicate 95% confidence intervals. (C) A three-way mixed ANOVA (Accuracy × Confidence × Group) on BOLD response in the TMS-vertex session, with individual data points superimposed on the bar plot. Error bars denote the standard error of the mean (SEM) over participants.

In order to fully characterize this BOLD-behavior relationship in the TMS-vertex session, we divided the volunteers into two subgroups using median split of meta-memory efficiency score (HIGH vs. LOW meta-memory efficiency subgroup; n = 9 each) and ran a three-way mixed ANOVA (Accuracy: Correct/Incorrect × Confidence: High/Low × Group: HIGH/LOW) on the individual BOLD response extracted with the aforementioned mask. We found a significant main effect of Confidence (*F* (1,16) = 8.93, *p* = 0.008) and a marginally significant three-way interaction (*F* (1,16) = 4.33, *p* = 0.054, Figure 3C), which was driven by a significant difference between high confidence vs. low confidence rating for correct trials in the HIGH meta-memory efficiency subgroup (*F* (1,8) = 10.14, *p* = 0.012) and its corresponding absence in the LOW meta-memory efficiency subgroup (*F* (1,8) = 0.16, *p* = 0.698). Following TMS administered to the precuneus, all these effects were not observed at all (all *ps* > 0.05). By taking individual variability in BOLD activation into account, the functional relevance of the precuneus in meta-memory efficiency is well evinced.

### Resting-state functional connectivity analysis (rs-fcMRI)

In addition to task-based BOLD responses, recent works have also identified single-neuron responses in the human posterior parietal cortex which appear to code recognition confidence [15], and suggested a stream that reads out meta-memory from the hippocampus in nonhuman primates [25, 33]. In order to measure information communication from distributed brain regions, we estimated the measure of functional integration between precuneus and hippocampus (HP distance) over resting-state BOLD response. To aid interpretation, the shorter the HP distance is, the stronger functional integration between precuneus and hippocampus (Figure 4A).

**Figure 4.**
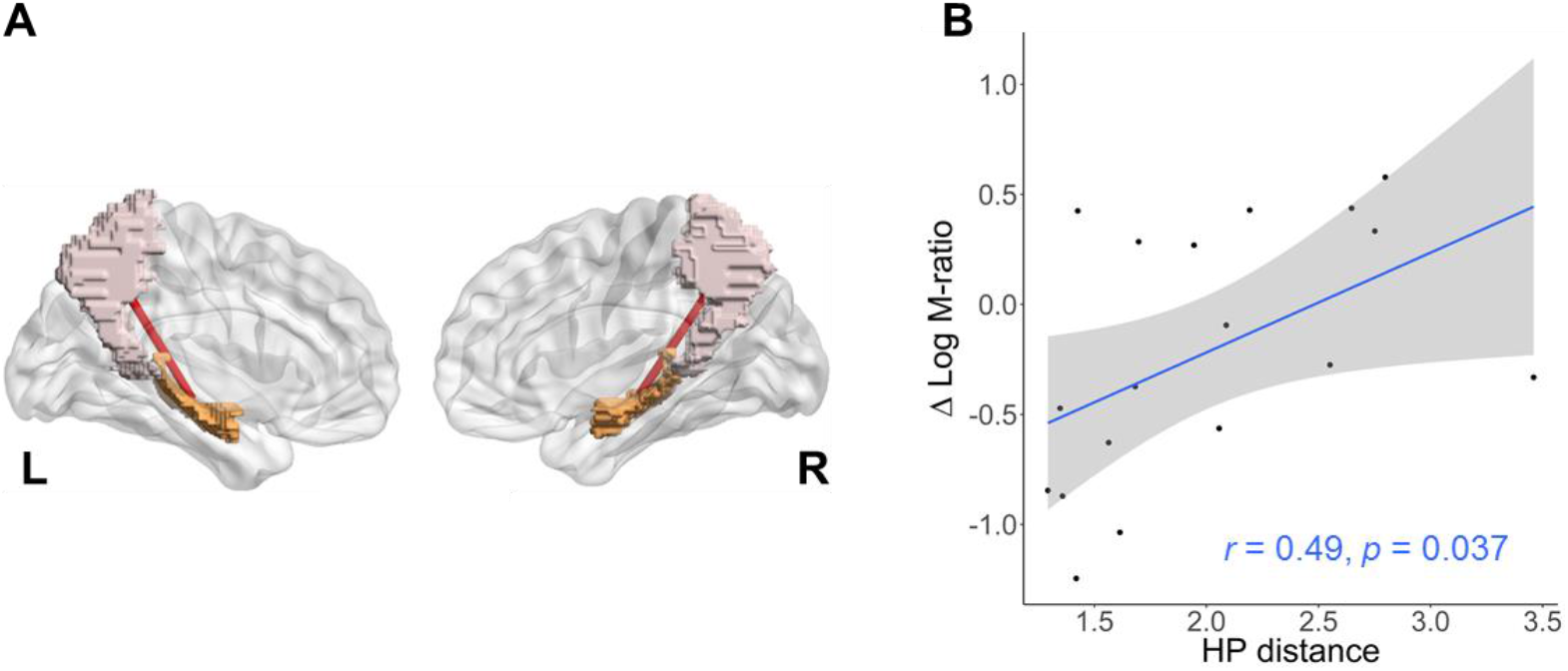
(A) A cartoon image to show hippocampus-precuneus (HP) connectivity distance between hippocampus (orange color) and precuneus (pink color) separately for left-and right-hemisphere. The resulting HP distance was averaged across hemispheres. (B) Scatter plot between HP distance and the change in metacognitive efficiency (TMS-precuneus > TMS-vertex), with 95% confidence intervals.

We investigated the effect of TMS on the association between HP distance and the change in meta-memory efficiency with a linear regression model, and a significant positive correlation was found (Pearson’s *r* = 0.49, *p* = 0.037, Figure 4B). Individuals with higher functional connectivity between precuneus and hippocampus showed higher vulnerability after TMS to the precuneus, compared to the vertex (TMS-precuneus > TMS-vertex). In order to show such effects to be specific to the memory domain, we also ran this correlational analysis with the change in metacognitive efficiency obtained from the perceptual task and found no relationship between HP distance and meta-perceptual efficiency (Pearson’s *r* =-0.36, *p* = 0.143); the two correlation coefficients were significantly different from each other (comparison between correlations: *z* = 3.27, *p* = 0.001).

### Voxel-based morphometry (VBM) analysis

Having observed that metacognitive ability was reduced by TMS to precuneus than to vertex, we then asked whether this inhibitory effect of TMS was predicted by variability in grey matter volume (GMV) in the precuneus across subjects. We investigated the association between GM volume and the change in meta-memory efficiency (TMS-precuneus > TMS-vertex) after controlling for total brain volume and gender (male/female). The results showed that change in meta-memory efficiency was positively correlated with GMV in the precuneus (*t* = 3.75, SVC-*P*fwe < 0.05 at x, y, z = 15, −68, 43; Figure 5A). We also ran the same analysis on the meta-scores obtained from the perceptual task and no association between precuneal GMV and change in meta-perceptual efficiency was found, again highlighting the domain-specificity of our main findings.

**Figure 5.**
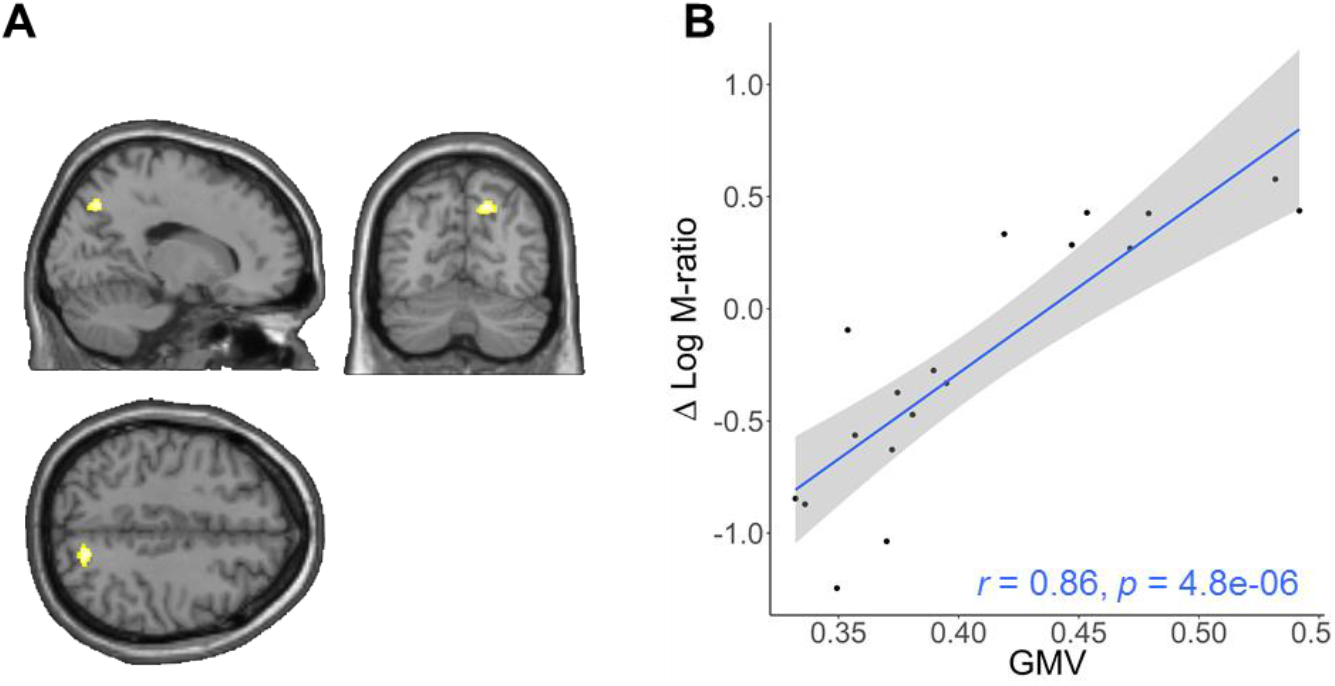
(A) Brain regions with positive correlation between grey matter volume (GMV) and difference in metacognitive efficiency (Log M-ratio in the TMS-precuneus session –Log M-ratio in the TMS-vertex session). The significant cluster was found in the precuneal region (*P*_FWE_ < 0.05 small volume correction). For display purpose, brain maps were thresholded at *p* < 0.005 uncorrected. (B) Scatter plot between individual GMV from the peak voxel (x, y, z = 15, −68, 43, right precuneus) and their change in metacognitive efficiency, with 95% confidence intervals.

To visualize this correlation pattern, we plotted the linear relationship between GMV and change in metacognitive efficiency scores at the peak voxel of this cluster across participants (Pearson’s *r* = 0.86, *p* < 0.001; Figure 5B). These results revealed that participants with a smaller volume/density in the precuneus tend to have higher vulnerability to TMS in metacognitive ability, whereas those with a bigger volume/density in this region tend to be more immune to the TMS disruption.

Finally, we ran a control analysis correlating individuals active motor threshold with his/her delta log M-ratio and showed that the putative effects by TMS on meta-memory ability was not modulated by the penetrability/thickness of the individuals skull per se (Pearson’s *r* = 0.13, *p* = 0.611), reinforcing our main findings that the neuromodulatory effects by TMS were specific to the precuneus-related anatomical profiles.

## Discussion

TMS on the precuneus was found to impair metacognitive efficiency in a long-term memory retrieval task without affecting type 1 task performance [18]. Despite reaching statistical significance, the effect size in that study was relatively small as in the effects were stronger in one measure than the other (i.e., fully statistically significant for Meta-d’ –’ but only marginally significant for Log M-ratio measure). The discrepancy between the two metrics might be potentially caused by the sizable individual differences among the participants, as of other reported observations that individual variability imposes sizable influences on determining the experimental effects of brain stimulation [34]. Here, we set out to quantify the neuromodulatory effects of TMS making use of individual differences in terms of BOLD responses and two anatomical profiles. Multimodal characterization as such illuminated unambiguously the importance of the precuneus in supporting meta-memory upon episodic recollection.

We first established that the precuneal region is functionally implicated in meta-memory judgement using task-related BOLD signal measurements. In contrast to the findings that BOLD activities in the right rostro-lateral prefrontal cortex are predictive of meta-perceptual ability [4], our findings showed an association between BOLD responses in the precuneus and meta-memory efficiency. Specifically the BOLD responses in the precuneus was correlated with individuals’ metacognitive efficiency in the control session, but such correlation was disrupted following TMS on the precuneus, pointing to a critical role of the precuneus in metacognitive ability for memory processes [3, 6, 8]. These results are in line with a recent report that meta-memory specific signals being located in the precuneus [6] as well as clinical findings that patients with lesions in the posterior parietal cortex tend to exhibit reduced confidence in their source recollection [16, 17].

A natural question to ask is to what extent and how the precuneus is mechanistically involved in meta-memory processing. While the overall BOLD level given by the interaction term was equated across the two TMS sessions at the group level, we showed that TMS to the precuneus considerably weakened the correlation between metacognitive efficiency and brain activation across subjects. This implies that the precuneus might be implicated to different extents across participants in subserving meta-memory assessment in face of the acute TMS disruption.

In affirmation of this notion, we revealed that individuals with higher functional connectivity between the precuneus and the hippocampus, or smaller GM volume/density in the precuneus, tend to exhibit higher vulnerability in metacognitive ability under the impact of TMS. By indexing the strength of functional connectivity between the precuneus and the hippocampus, we showed that subjects with higher functional connectivity were more vulnerable to the inhibitory TMS effect. Since the hippocampus is crucial for temporal order memory judgement [35] and is known to modulate the neural activity of confidence judgments [19], we propose that the precuneus might act as an accumulator [36] for the strength of evidence received from hippocampus, which was also utilized to support meta-mnemonic/meta-awareness appraisal. Although how the information is concurrently transformed for meta-memory processing is still unknown, our results indicate that this “meta-mnemonic” accumulator during memory retrieval was dependent on its functional connectivity with the hippocampus. In fact, neuroimaging with pharmacological intervention on the monkeys has delineated a meta-memory stream consisting of information flow extracted from the hippocampus, going through the intraparietal cortex and then read-out by the prefrontal area 9 [25, 33]. Using high-resolution multi-parameter mapping, researchers also found that markers of myelination and iron content in the hippocampus correlate with metacognitive ability across individuals [2]. Altogether, these may help account for the neuromodulatory effects by TMS being dependent on individuals’ functional connectivity between the precuneus and the hippocampus.

We further revealed that the changes in metacognitive efficiency following TMS were determined by the GM volume/density in the precuneus. Specifically, our participants with a smaller volume/density in precuneus tend to have higher vulnerability in metacognitive ability to TMS, whereas those with a bigger volume/density in this region tend to be more immune to the TMS impact. The correlational relationship between precuneal volume and meta-memory capability has been previously established during a verbal memory task [3]. Here, in light of the findings that the posterior parietal cortex contains two sub-groups of neurons that are differentially responsive for memory versus confidence demands during memory retrieval

[15], our revelation that the precuneal density/volume is a robust predictor for individuals’ susceptibility might thus align with the possibility that participants with a bigger/denser precuneus might have a larger “missed” portion of the precuneus that can remain functional to serve to faithfully code the confidence-related signals.

## Conclusions

Taken both functional and anatomical evidence together, our study capitalized on individual variability to characterize the neuromodulatory effects of TMS during meta-mnemonic appraisal. Through several neuroimaging modalities, we provided compelling evidence in outlining a possible circuit encompassing the precuneus and its mnemonic midbrain neighbor the hippocampus at the service of realizing our meta-awareness upon memory recollection of episodic details.

## Acknowledgements

We thank Elena Makovac for her advice on implementing VBM analysis. The authors declare no competing financial interests.

